# Unequal barriers to nature’s contributions to people impact quality of life

**DOI:** 10.64898/2026.01.19.699912

**Authors:** M. Neyret, S. Lavorel, B. Locatelli, B. Martín-López

## Abstract

While nature contributes to multiple dimensions of human quality of life, access to nature and its benefits is unequally distributed. We surveyed residents of a French metropolitan area to address three questions: (i) what are the most common barriers to accessing various Nature’s Contributions to People (NCP)? (ii) how do these barriers vary across socio-demographic groups? (iii) how do they relate to nature connectedness and impact quality of life? Most frequent barriers, i.e. low availability or quality of green spaces, lack of time, and crowding, differed across NCP categories. Individuals with lower income, limited physical mobility, living in urban areas, and women reported more barriers. More numerous barriers reduced NCP satisfaction, with repercussions on quality of life. In contrast, nature connectedness directly increased quality of life. Our finding that barriers accumulate for disadvantaged groups, reinforcing social inequities, calls for policies addressing multiple dimensions of access to promote equitable relationships with nature.

**Graphical Abstract:** Men, people living in rural areas and those with higher income or mobility encounter fewer barriers to accessing and benefiting from Nature’s Contributions to People (NCP), and thus report greater satisfaction with NCP. They also report stronger nature connectedness. Both pathways, in addition to direct effects, lead to higher reported quality of life for these individuals.

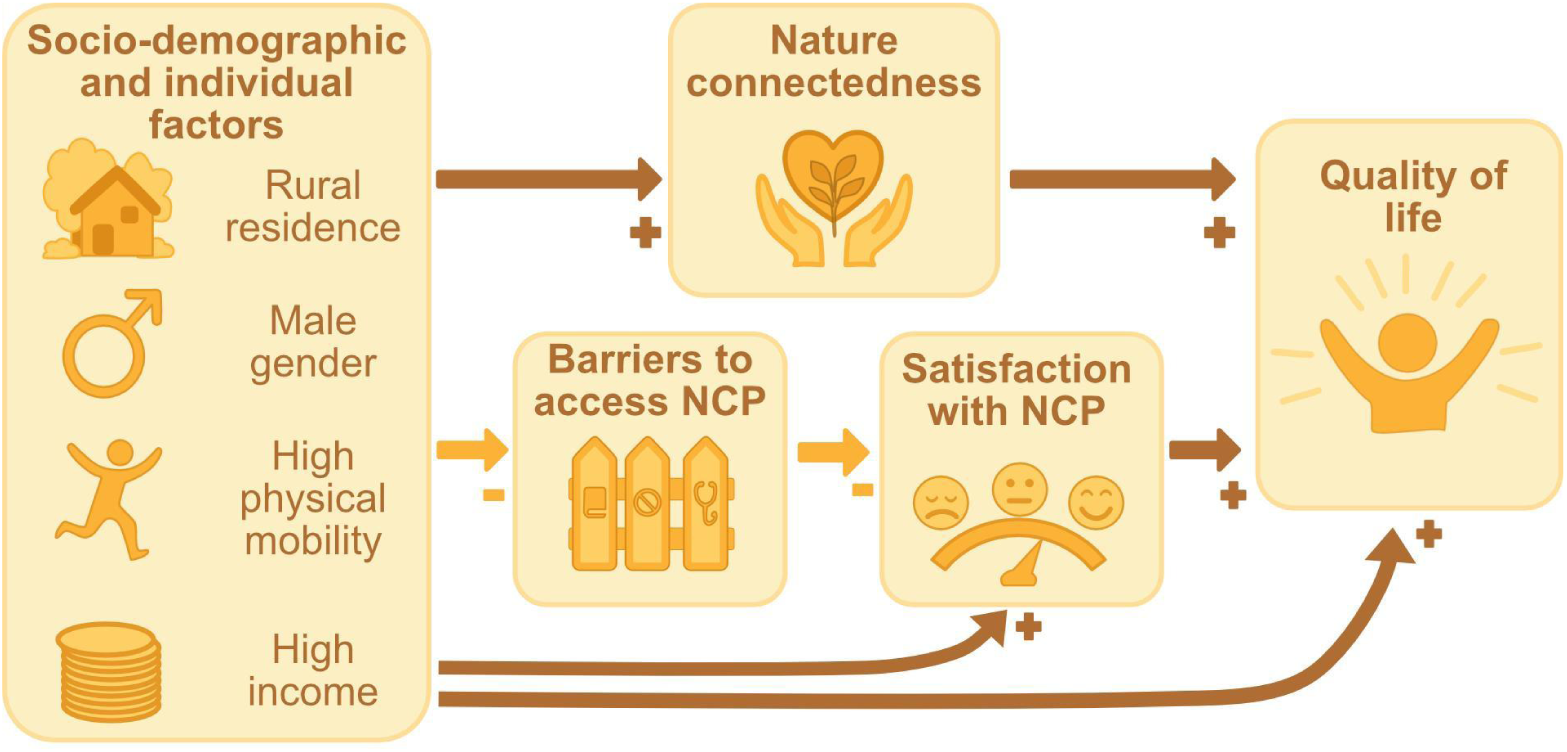

## Introduction

Nature contributes to many dimensions of human well-being. It supports subsistence through food and timber production, provides protection from natural hazards such as floods and erosion, and maintains air and water quality (Díaz et al. 2015). Green spaces frequentation also enhances mental and physical health as well as social relationships (Aerts et al. 2020; Marselle et al. 2020). Managing ecosystems to increase these contributions to people - for example, by expanding urban green areas (Reyes-Riveros et al. 2021), allowing access to forest or shellfish resources (Chaudhary et al. 2018; Wieland et al. 2016) - has often been assumed to benefit society as a whole. However, such aggregated approaches overlook the social heterogeneity that shapes people’s ability to benefit from nature, thereby concealing the existence of winners and losers (Daw et al. 2011). Indeed, differences in priorities, needs, and power - linked to wealth, ethnicity, physical ability, and other factors - produce disparities in who actually benefits (Locatelli et al. 2025). For instance, Chaudhary et al. (2018) documented unequal access to forest resources in Nepal based on caste, income, wealth, and gender; and Adekola et al. (2015) found that the distribution of environmental costs and benefits has strongly disadvantaged local communities in Nigerian wetlands. Studies in the global North have also shown that ethnic minorities and people with disabilities benefit less from green spaces (Edwards et al. 2023; Morris et al. 2011; Winkler et al. 2025).

Considering the essential role of nature’s contributions to people (NCP) in supporting the material, mental, and social dimensions of quality of life (Ausseil et al. 2022; Lapointe et al. 2020; Martín-López et al. 2018; McCartan et al. 2023), unequal access to these benefits is likely to generate disparities in well-being and may further disadvantage already marginalised populations (Aznarez et al. 2024; Gourevitch et al. 2021). For example, greater barriers to accessing outdoor spaces have been linked to lower reported quality of life among older adults (Rantakokko et al. 2010). Because access barriers limit or prevent nature experiences, they may also increase nature disconnection (Beery et al. 2023). Nature connectedness - the extent to which individuals feel emotionally, cognitively, and physically part of the natural world (Schultz et al. 2002; Nisbet et al. 2009) - has been associated with better mental health, vitality, and overall well-being (Capaldi et al. 2014; Zeng et al. 2025). It also strengthens the impact of nature contact on well-being (Martin et al. 2020). Yet nature disconnection is increasing worldwide, driven by factors such as urbanisation, environmental degradation, and shifts in lifestyle and education (Soga and Gaston 2023), and has itself been linked to poorer mental health and quality of life (Martyn and Brymer 2016; Zeng et al. 2025). Access barriers may thus contribute to a reinforcing cycle: reduced nature contact and associated “extinction of experience” weaken connectedness, which in turn generates additional psychosocial barriers - such as fear of the unfamiliar - further diminishing contact with nature (Soga and Gaston 2016). Understanding what drives unequal access is therefore crucial for developing inclusive environmental policies that promote nature contact and improve quality of life across diverse populations.

Several frameworks conceptualize access to nature from an environmental justice perspective. Focusing on poverty alleviation, Sen’s capability approach (Sen 1999) emphasizes that well-being depends not only on resources but also on people’s capabilities, such as health, location, and social circumstances. This approach has, for instance, been applied to outline individual and collective ability to access NCP in the Alps (Grosinger et al. 2021). Ribot and Peluso (2003) defined access as “the ability to derive benefit from things,” highlighting the rights-based processes that shape access to resources. Building on these ideas, Szaboova et al. (2020) identified four types of mechanisms shaping access to green spaces: rights-based (use and control rights), physical (factors that facilitate, or hinder, people’s ability to physically access nature, such as transport), structural and relational (e.g. social networks), and psychosocial (e.g. preferences, or perceptions of safety).

Despite this recognition of access as multidimensional, there remains little synthesis of the barriers people face when accessing a wide range of NCP, or of how such barriers vary across social groups. Much of the literature typically focuses on identifying barriers to access natural spaces without consideration of NCP (Arnberger et al. 2017; Edwards et al. 2023; Morris et al. 2011; Rigolon 2017; Winkler et al. 2025) or considering a restricted set of NCP (Lau et al. 2020; Wieland et al. 2016). These studies also tend to focus on specific groups, such as ethnic minorities (Edwards et al. 2023; Morris et al. 2011) or age groups (Arnberger et al. 2017; Rigolon 2017). As a result, quantitative comparisons of the barriers encountered across multiple NCP and socio-demographic groups remain scarce (but see Lapointe et al. 2020).

Drawing from the fields of environmental justice, ecosystem service, and welfare studies, this paper aims to fill two research gaps: (i) the lack of quantitative comparison of the prevalence of nature barriers across social groups; and (ii) our poor understanding of how these barriers interact with nature connectedness and impact quality of life. More specifically, we ask three questions: (i) What are the most common barriers to access NCP and do their prevalence vary across socio-demographic groups? (ii) How do these barriers affect people’s satisfaction with the NCP they benefit from? and (iii) How do these barriers interact with nature connectedness and impact people’s quality of life? Our hypothesized framework is shown in Fig. 1. Specifically, we hypothesized that urban residents, people with lower physical mobility, lower income, and women are more likely to report a higher number of barriers. Further, we assumed that a high number of barriers would be associated with low satisfaction with highly valued NCP and low reported quality of life, when accounting for other socio-demographic variables. We tested these hypotheses using an online survey with more than 700 responses from the general public in the ecologically and socially diverse area of Grenoble in South-Eastern France.

**Figure 1.**
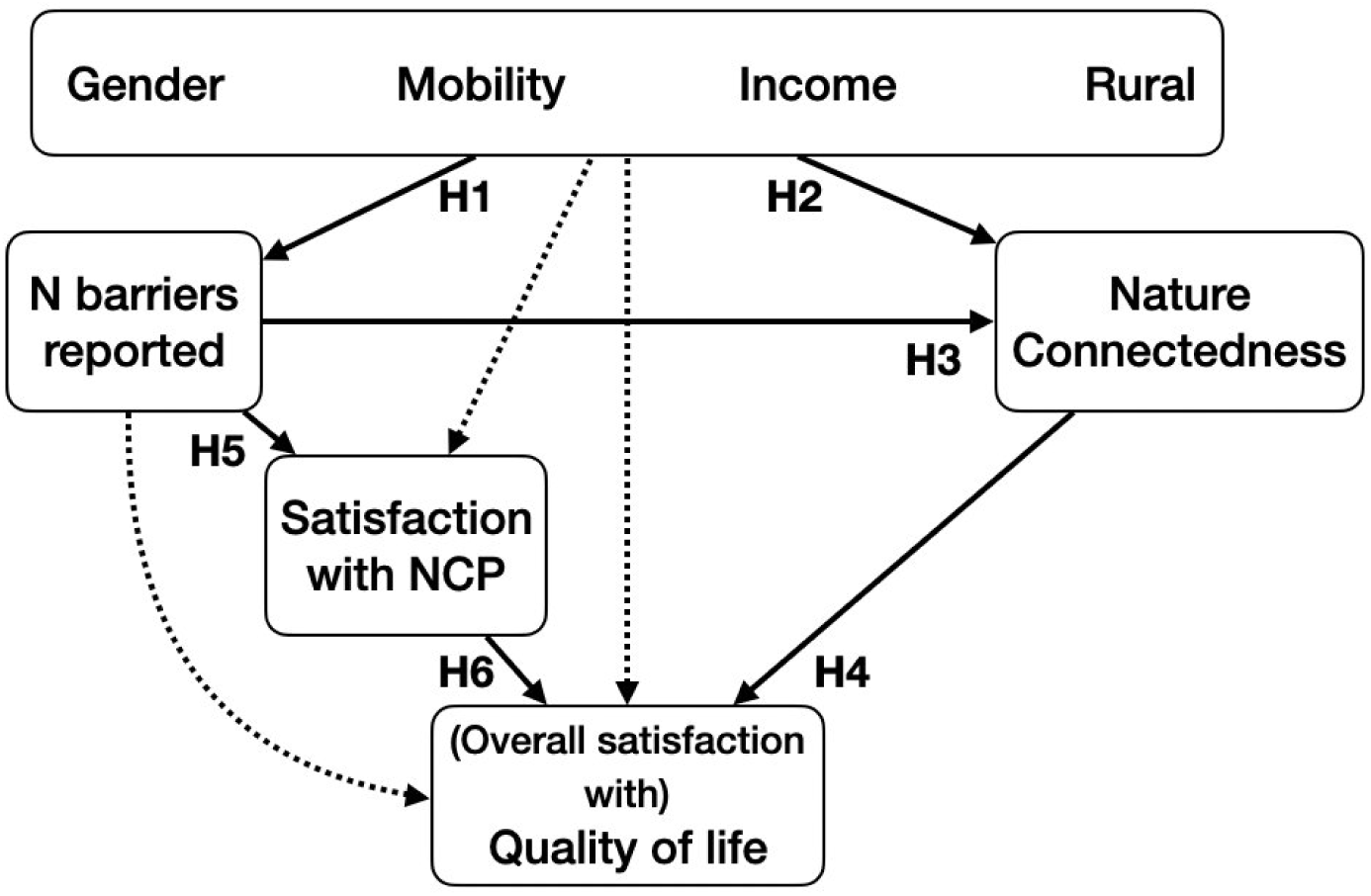
Structure of the hypothetical framework linking socio-demographic factors, access barriers, nature connectedness, satisfaction with NCP and quality of life. Socio-demographic factors are represented within a single box but were considered independently. The structure was built based on the following hypotheses. **H1.** Women, disadvantaged groups and those living in urban areas face more barriers in accessing NCP (Lapointe et al. 2020; Winkler et al. 2025; Morris et al. 2011). **H2**. Socio-demographic variables, such as physical mobility and place of residence, impact nature connectedness (Macias-Zambrano et al. 2024). **H3**. More barriers in accessing NCP lead to lower exposure, resulting in lower nature connectedness (Beery et al. 2023; Zeng et al. 2025). **H4**. Nature connectedness is positively associated with quality of life (Zeng et al. 2025). **H5 and H6.** Barriers to access important NCP affect satisfaction with NCP and quality of life. Dotted arrows represent paths that did not have a specific hypothesis of interest to this study associated with; they were included to control for potential additional covariations (for instance, a positive effect of income on quality of life independently from NCP and nature connectedness).

## Methods

### Study region

The study region includes the urban, peri-urban, rural and mountainous surroundings of the Grenoble metropolitan area (45°10’ N, 5°43’ E) and surrounding intermunicipalities, in South-Eastern France. It spans 4450 km², including 479 municipalities and a total of approximately 950 000 inhabitants. The area is structured by three biodiverse mountain ranges - Vercors, Chartreuse and Belledonne - which benefit from a wide range of protection measures through two natural parks and several conservation areas. These mountains delineate three valleys (Drac, Isère and Bièvre) that host agricultural activities but are also under pressure from urban sprawl (Desgouttes and Gilbert 2014; Vannier et al. 2016). The region thus faces tensions between nature conservation, attractiveness for residents and tourists, and sustainable agricultural production (Vannier et al. 2019).

Most of the population in the area is concentrated in the urban Grenoble metropolis. Socio-economically, the area has a higher proportion of white-collar and intellectual professions, and fewer workers involved in industry, compared to the national average (Desgouttes and Gilbert 2014). Although the share of low-income households is lower than in other French metropolises, disparities remain stark, with most low-income households (including many students with no or limited direct income) concentrated in the south of the metropolis, and much higher average incomes in the north-east (Decorme and Labosse 2021). Our study area also encompassed surrounding rural and mountain areas, with economic activities centered on cropping, low- to medium-intensity livestock farming, and tourism.

### Data acquisition

#### Survey design and distribution

We designed and conducted an online survey between March and October 2024 using the Maptionnaire software (see detailed survey in the supplementary material). The survey was pre-tested in two rounds of 10-15 respondents each to assess user-friendliness and clarity, and adapted accordingly. It was approved by the first author’s university (Université Grenoble Alpes) data protection officer before distribution.

To circulate the survey, we reached out to local administrations (Grenoble metropolitan area, regional planning, water and environmental agencies, regional parks, inter-municipal cooperation bodies, the Chamber of Agriculture, and municipalities), as well as environmental NGOs, associations (hiking, fishing, hunting), and academic institutions. These contacts were asked to share the survey both internally and with their broader audiences. In addition, leaflets and posters were distributed in shops and cultural centers. Consequently, we did not aim for, nor obtained, a representative sample of the whole population, and our sample is likely biased towards people with high education and interest for nature.

Respondents were first requested to locate their residence and assess their overall quality of life (self-reported quality of life: “I am satisfied with my quality of life and well-being” - 4-point Likert scale). Then, they had to choose the three NCP most important to their well-being from a list of fifteen NCP, selected from previous work as the most valued NCP in the area (Vannier et al., 2019). These included two material NCP (crop and timber production), six regulating NCP (temperature cooling, protection against floods and against erosion by vegetation, regulation of air and water quality, and global climate regulation) and seven non-material NCP (outdoor recreation in urban or peri-urban settings, outdoor recreation in the countryside or mountains, cultural identity, habitats for biodiversity, aesthetic enjoyment, wild product harvesting, and hunting/fishing). Habitats for biodiversity were considered as non-material as we assumed most respondents would relate to the cultural and intrinsic value of biodiversity. Harvesting and hunting/fishing were also considered as non-material NCP because they are locally considered as hobbies rather than food sources. These decisions regarding NCP classification, while appropriate for our study, should be adapted for each study as they may vary across socio-cultural contexts; however, they are unlikely to impact our main results on links between barriers, connectedness and quality of life, which do not rely on NCP categories.

For each chosen NCP, we asked respondents to rate their satisfaction with their ability to benefit from each NCP (four-point Likert scale) and to report barriers limiting this ability. Participants could select any number of barriers relevant to them from a list of barriers provided in the questionnaire (Table 1), or suggest additional barriers. Finally, respondents were asked about their socio-demographic and individual background, including gender, age, education, income, physical mobility, length of residence in the study region, childhood setting (in or outside the region, rural or urban), nature connectedness using the inclusion of Nature in Self scale (Martin and Czellar 2016), and weekly time spent in nature. Our survey did not integrate data on race or ethnicity due to ethics clearance.

**Table 1.** Barriers listed for each Nature’s Contribution to People (NCP). Non-material NCP include outdoor recreation in urban or peri-urban settings, outdoor recreation in the countryside or mountains, cultural identity, aesthetic enjoyment, hunting and fishing, harvesting of wild foods. Material NCP include crop and feed production, timber production. Other regulating NCP include protection against floods, global climate regulation, air quality, water quality, protection against erosion.

| Barrier to access NCP | Description in the survey | Non-material NCP | Material NCP | Regulating NCP |  |
| --- | --- | --- | --- | --- | --- |
|  |  |  |  | Temperature cooling | Other regulating NCP |
| Low availability (of green or natural spaces providing the NCP) | There are no or insufficient green or natural spaces nearby that provide this benefit. | X | X | X | X |
| Low quality (of green or natural spaces providing the NCP) | The spaces that provide this benefit are not of sufficient quality or are in poor condition (dirty, damaged, etc.). | X | X | X | X |
| Lack of knowledge | I don't know where to go or I don't know which spaces can provide me with this benefit. | X | X | X | X |
| Cost | It is too expensive. | X | X | X |  |
| Legal | I am not legally allowed to enjoy this benefit where I would like to (private area with restricted access, protected space, etc.). | X | X | X |  |
| Lack of time | I don't have time, I'm too busy. | X | X | X |  |
| Crowding | There are too many other people, or other people annoy me and prevent me from enjoying this benefit (noise, commotion, etc.). | X |  | X |  |
| Lack of transport | I cannot easily access the spaces that provide this benefit due to a lack of roads or public transport. | X |  | X |  |
| Lack of companionship | I have no one to go with to the spaces that provide this benefit. | X |  | X |  |
| Belonging | I don't feel comfortable because I feel isolated or marginalised in the spaces that provide this benefit. | X |  | X |  |
| Culture | I don't feel comfortable because the spaces that provide this benefit are not suited to my beliefs or culture. | X |  | X |  |
| Lack of infrastructure | There is not enough infrastructure (benches, toilets, signs, etc.) in the spaces that provide this benefit. | X |  | X |  |
| Lack of motivation | I lack motivation. | X |  | X |  |
| Safety (social) | I do not feel safe in these spaces because of other people. | X |  | X |  |
| Safety (environmental) | I do not feel safe in these spaces because of the environment (fear of getting lost, falling, being bitten by ticks or insects, etc.). | X |  | X |  |
| Poor health | My health does not allow me to access or enjoy these spaces (physical condition, pain, allergies, asthma...) | X |  | X |  |

#### Identification of barriers limiting the ability to benefit from NCP

Building on barriers reported in the literature (Edwards et al. 2023; Lapointe et al. 2020; Morris et al. 2011; Sefcik et al. 2019), adapted to our local context, we created a list of 16 barriers that encompass the physical, legal, structural/relational and psychosocial dimensions of access to nature (Szaboova et al. 2020). Many of those barriers were not relevant for all NCP (e.g., lack of motivation for air quality regulation), and including all as options for each NCP created confusion among survey testers. Hence, we decided to pre-select which barriers would be offered for each NCP category. The barriers were identical within each NCP category (material/non-material/regulating) except for temperature regulation, a regulating NCP that partly requires physical access to green spaces and thus shared all barriers with the non-material NCP. Table 1 shows the list of all barriers limiting the ability to benefit from an NCP (hereafter referred to as barriers), their corresponding prompts in the survey, and to which NCP they were applied.

However, respondents could also report additional barriers as free-text responses, which were reclassified into existing categories or clustered into new ones, resulting in additional barrier categories - such as sustainability, related for instance to the reluctance to use a car to access mountain areas. Barriers associated with noise and air pollution related to cars, industry, or pesticides were classified as a lack of quality of the green space. Safety barriers due to the presence of hunters, other users’ pet dogs or livestock protection dogs were classified as social insecurity (hunters and pet dogs) and environmental insecurity (protection dogs).

Importantly, this means that barriers were not measured, but self-reported. Differences in perception or willingness to report a barrier might thus contribute to observed differences, although perceived barriers can arguably be as effective as objective ones in limiting access (Yasumoto et al. 2021).

#### Data analysis

All the analyses were run using R. Key packages included ordinal, lavaan, glmmTMB. Visual representations relied on the ggplot2 and ggthemr packages.

#### Sample description

A total of 711 respondents from 240 municipalities completed the survey. Respondents were relatively balanced in terms of gender, age (between 18 and 74 years, with few respondents older than 75 years), rural or urban/peri-urban residence, and childhood in or outside the study region (Fig. S1). 67% of respondents grew up in the countryside and mountain areas rather than in urban areas. Missing values represented 13% of all responses for income, < 2% for childhood residence and diploma and < 1% for all other variables.

We observed exceptionally high engagement among members of the regional hunting association, which led to an over-representation of hunters in our survey compared to the regional statistics. Based on reported priorities and activities, 199 respondents out of 711 were assumed to be closely associated with hunting or fishing activities (∼28%) - while the county’s hunting association counts around 16 000 licensees for around 950 000 adult inhabitants (i.e. 1.7%). To maintain representativeness in our results, we applied a weight of 10% - a round number accounting for the fact that some respondents classified as potential hunters may in fact be fishers (as both activities were grouped together) - which brought their effective contribution to 3%, slightly above the regional proportion. This weighting had little impact on the overall results: an equivalent unweighted SEM (see below and suplementary material) yielded similar coefficients, though with a significant p-value, suggesting a poorer fit to the data, likely due to the larger effective sample size rather than a substantive difference in model structure.

Socio-demographic variables and Likert scales were converted into ordinal, semi-quantitative scales (e.g., from 1 to 4). We also associated respondents’ place of residence with a degree of rurality based on the Global Human Settlement index from 7 (Very Low Density Rural grid cell) to 1 (Urban Center grid cell). We then conducted a Principal Component Analysis (PCA) on those variables (age, income, time spent in the region, childhood environment, childhood in the region, gender, education, physical mobility, and degree of rurality of the residence) to identify the main axes of variation in our sample and select key socio-demographic factors (Figure S2). The main axes of variation among our respondents were: i) a gradient from urban residents who spent little time in nature and had low nature connectedness, to rural residents with higher nature connectedness, who spent more time in nature, and typically had lived in the region for longer (23.9% variance); ii) a gradient from low to high income, which was also associated with childhood environment (16.4%), iii) physical mobility (13.8%) and iv) gender (11.1%). Consequently, to cover the spread of socio-demographic and individual factors in our sample, we selected place of residence (rural to urban gradient), income, physical mobility and gender as focus variables for our analysis. These factors correspond to drivers of environmental disparities commonly identified in the literature (Lapointe et al. 2020; Winkler et al. 2025).

#### Analysis of barrier prevalence across groups

We calculated the frequency of each barrier within each NCP category. Then, we assessed how the number of barriers and the chance of selecting each individual barrier varied across socio-demographic variables. We fitted generalized linear mixed models for each barrier (or number of barriers) as the response variable and all socio-demographic variables (function glmmTMB, package glmmTMB). For non-material and regulating NCP, we included random effects for NCP to account for non-independence. There were only two material NCP, so we could not include them as random effects and used fixed effects instead. As an alternative approach, we fitted models with pairwise interactions between socio-demographic variables; most interactions were weak, and these results are not presented. We did not correct p-values for multiple comparisons because each model addressed a distinct outcome (a specific reported barrier), and the explanatory variables were selected *a priori* as part of our single theoretical framework rather than through exploratory screening.

Finally, we assessed whether barriers to access NCP impacted respondents’ satisfaction with the NCP provided. We tested the relationship between the number of barriers reported by respondents and their reported satisfaction with the corresponding NCP. Satisfaction was an ordinal factor with four levels from very low (1) to very high (4) satisfaction. We thus applied cumulative link mixed models with satisfaction as the response variable and the number of reported barriers as the explanatory variable, separately for e*ach NCP category.* NCP were included as random effects for regulating and non-material NCP.

#### Linking barriers with socio-demographic variables, nature connectedness and quality of life

We then tested the relationships between socio-demographic variables, nature connectedness and quality of life. As nature connectedness and quality of life were ordinal factors, we applied cumulative link mixed models with socio-demographic variables as explanatory variables. NCP were included as random effects.

Finally, we aimed to assess the relationships between socio-demographic variables and quality of life, the number of barriers reported, satisfaction with NCP, and nature connectedness. The structure of the hypothetical framework and the underlying hypotheses are detailed in Fig. 1. Because we aimed to include mediating factors and assess these relationships simultaneously, we fitted a Structural Equation Model (SEM) rather than a set of regressions. This approach has been used in other studies to investigate the inter-relations between ecosystem services and well-being (Aldana-Domínguez et al. 2022). We used the lavaan package. Because of its limitations (i.e., assuming linearity and without random effects), these results were checked against a piecewise SEM, yielding similar results. Model details and results are presented in the supplementary material.

## Results

Most respondents (67%) reported at least one barrier to access NCP. Across all respondents and NCP, the most commonly reported barriers were the low availability of green spaces providing the NCP (17.4% of all reported barriers), lack of time (14%), and crowding preventing enjoyment of these spaces (13.7%). The main barriers differed across NCP categories, but were somewhat consistent within categories with some exceptions. Detailed results are shown in the supplementary material (Fig. S3, S4).

### Marginalised groups tend to report more barriers to access NCP

The number of barriers reported differed across socio-demographic groups (Table S1). High-income, physically mobile people and rural residents reported fewer barriers for non-material NCP. Men and rural residents reported fewer barriers for regulating NCP (Fig. 2).

**Figure 2.**
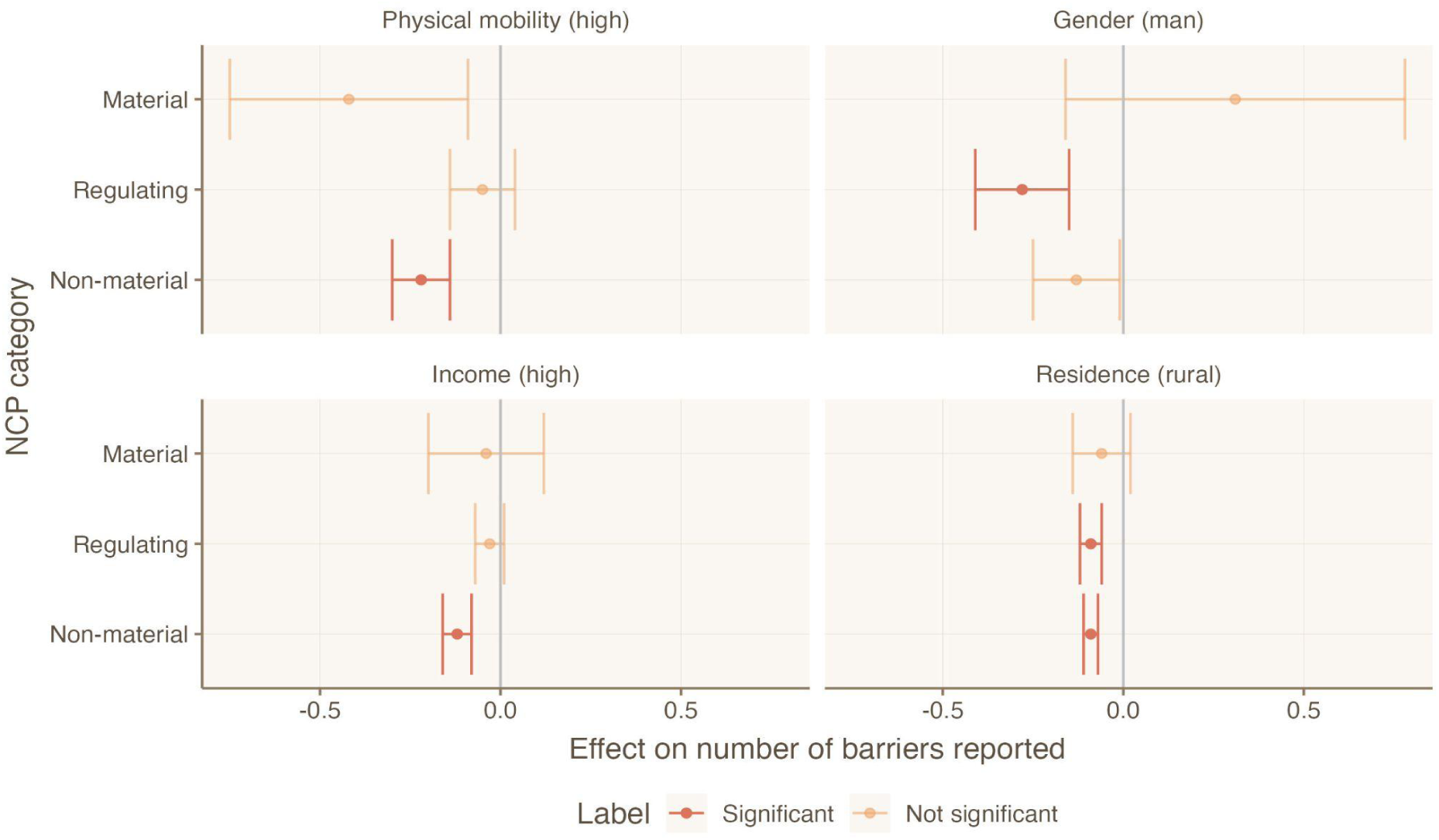
Number of barriers reported according to different socio-demographic factors. Bars show the model estimate (+/- standard error) of the socio-demographic factors’ effects on the number of barriers reported for each category of NCP. Red colour indicates that the response is significant (P < 0.05). If the estimate (central dot) is on the left of the vertical line, it indicates fewer barriers for people sharing the corresponding factor (rural residence, high income, male gender, high physical mobility). Results are shown for mixed models with the number of barriers as a response variable, and all socio-demographic factors as explanatory variables, and fitted independently for each NCP category.

The type of barriers reported also differed among groups (Fig. 3; see Table S2 for model parameters and p-values). People with good physical mobility were much less likely to report barriers related to poor health. They also reported fewer barriers related to lack of motivation. Men were less likely to report barriers related to poor health, lack of transports and knowledge. People with high income were much less likely to report cost-related barriers; this was also the case for barriers related to low quality of green or natural spaces. Although we were not able to test it directly, this is likely due to differences in lifestyle and place of residence, as they might benefit from higher-quality green spaces nearby or have more ease of transport, for instance due to car ownership. Finally, rural residents were less likely to report transport-related barriers than their urban counterparts, as they live closer to natural areas. They were also less likely to report barriers related to lack of motivation, time, knowledge, and availability (Fig. 3).

**Figure 3.**
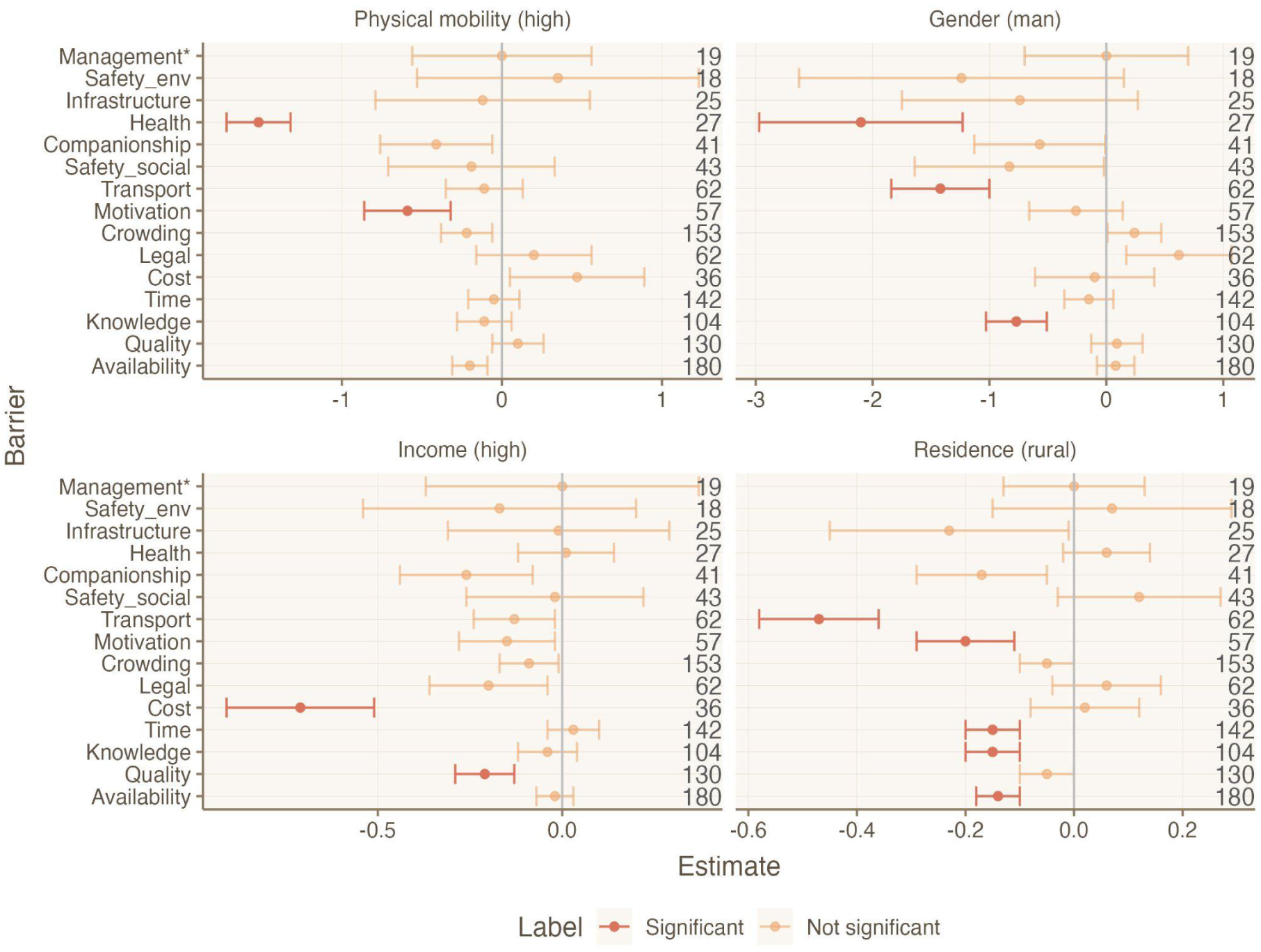
Chance of reporting of individual barriers depending on socio-demographic factors. Bars show model estimate (+/- standard error). Red colour indicates that the response is significant (P < 0.05). If the estimate (central dot) is on the left of the vertical line, it indicates fewer barriers for people sharing the corresponding factor (rural residence, high income, male gender, high physical mobility). Results are shown for mixed models with barrier (coded as binary: reported or not reported) as a response variable, and all socio-demographic factors as explanatory variables. Numbers on the right indicate how many times each barrier was reported; only barriers which were selected by more than 15 respondents are shown. Barriers marked * were not offered as options but added as free text by the respondents.

### Access barriers impact satisfaction with NCP and quality of life

Most respondents were satisfied with their ability to access and benefit from NCP (77% strongly or mostly agreed with the statement “I am satisfied with my ability to access and benefit from this NCP”); respondents from rural areas and with higher physical mobility tended to report higher satisfaction (Table S3). However, people who reported more barriers for a given NCP were less likely to be satisfied with their ability to access and benefit from this NCP (P < 0.001), suggesting a correlation between barriers and ability to benefit from NCP.

Rural respondents, men and physically mobile respondents reported higher nature connectedness (Table S3). Respondents living in rural areas, with higher physical mobility, and higher income reported higher quality of life (Table S3). As indicated by the structural equation model (Fig. 4), these effects were mediated by lower number of barriers for these groups, which was associated with higher satisfaction with NCP and quality of life, and by high nature connectedness, which directly and positively affected quality of life.

**Figure 4.**
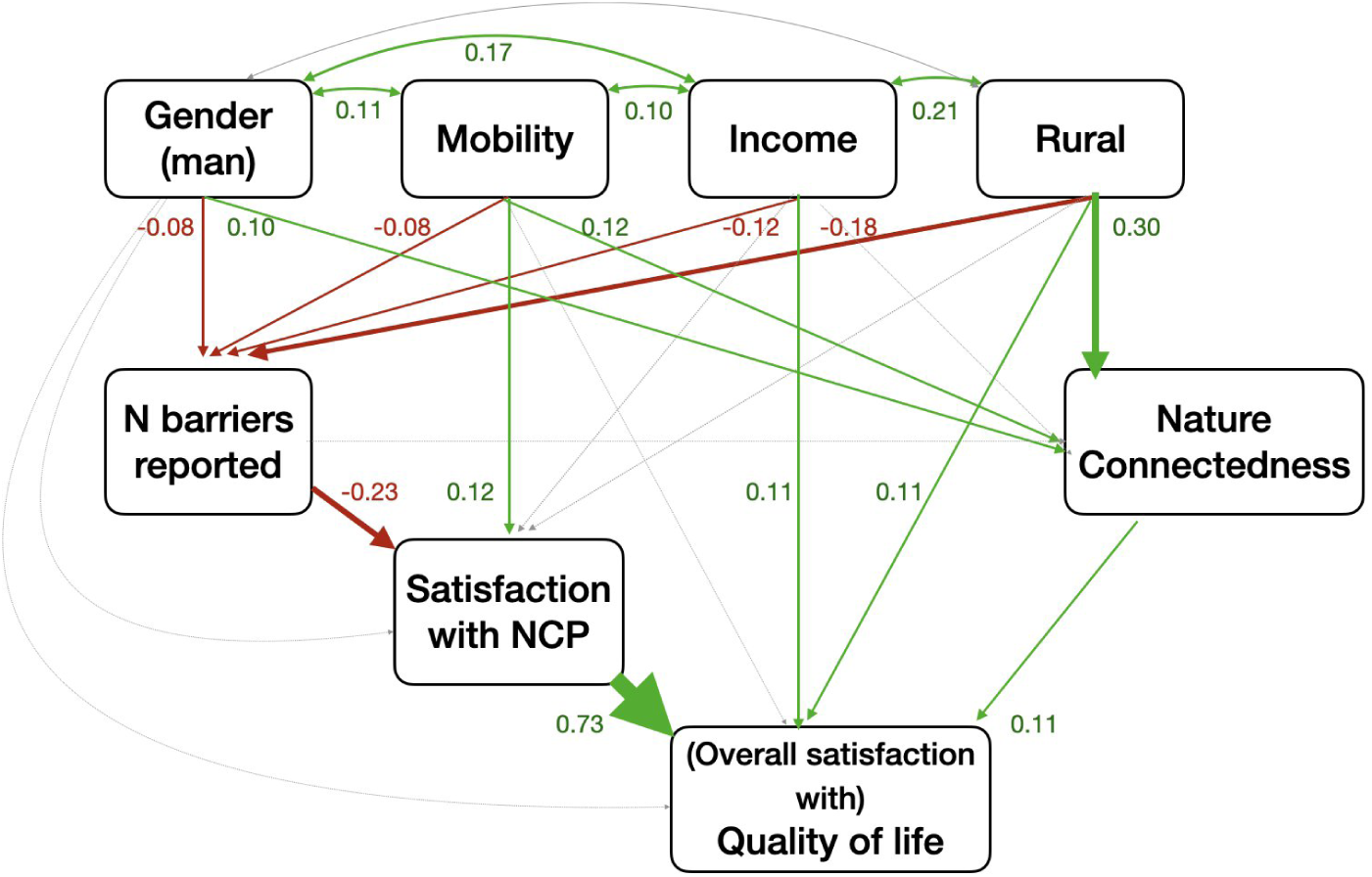
Structural equation model showing the mediating effect of access barriers, satisfaction with NCP and nature connectedness on quality of life. Red arrows indicate negative effects, green arrows indicate positive effects, grey arrows indicate non-significant paths. Numbers show standardised estimates. Bidirectional arrows show covariance between two variables.

## Discussion

In this study, we aimed to assess disparities in the barriers encountered by different groups to access and benefit from NCP, and the repercussions on their quality of life. Many respondents reported few barriers, and were overall satisfied with their ability to access and benefit from NCP. Yet, we highlighted that some groups tended to report more barriers, which was associated with lower satisfaction with the NCP provided; they also reported lower nature connectedness, with overall negative repercussions on quality of life.

### Accumulation of barriers for marginalised groups

We identified contrasting patterns of barriers across NCP categories and socio-demographic groups. Common barriers were often closely tied to characteristics of the NCP-providing spaces themselves, such as limited availability and poor environmental quality. Psychosocial and relational mechanisms-related barriers (Szaboova et al., 2020) were also prominent, in particular the lack of time available to enjoy nature. The presence of other users (crowding) was also a major deterrent to people’s ability to benefit from NCP. This matches well-known overcrowding issues of green and natural areas in the study region, and respondents perceived it as a direct (e.g., overuse of urban green spaces, presence of other users constraining hunting opportunities) and indirect limitation, through degradation that affects biodiversity, cultural identity, and aesthetic value. Overall, our results likely undercount barriers, as we only asked about the three NCP participants deemed most important. Respondents may thus have prioritised NCP that already contribute substantially to their quality of life and involve fewer barriers. Besides, pre-selecting barriers might have obscured barriers that we overlooked during the survey design.

Barriers were unevenly distributed: individuals with lower income, those living in urban settings, and those with limited physical mobility reported more barriers to access NCP, and lower associated satisfaction. This supports our hypotheses H1 and H5 (Fig. 1) and is consistent with previous studies that showed that urban and vulnerable groups as well as women have lower access to nature and the benefits it provides. For example, lower availability of high-quality urban green spaces - and the NCP they support - has been documented for low-income or minority households in France (Liotta et al. 2020), Germany (Kabisch and Haase 2014), Austria (Neier 2023), and several Central and Eastern European cities (Kronenberg et al. 2020). Beyond availability and quality, our work and others’ (e.g. Edwards et al., 2023; Rigolon, 2017) emphasise the importance of additional barriers that differentially shape people’s ability to benefit from NCP. These include governance and rule-related constraints, as well as individual capabilities and resources (Locatelli et al., 2025) across NCP categories. Examples include entry fees to urban parks reducing low-income residents’ access to material NCP in India (Basu and Nagendra 2020) and weak social networks and low perceived health diminishing older adults’ visitation of green spaces in Germany (Enssle and Kabisch 2020). Such barriers can lower perceived availability - and thus access to nature - even where “objective” accessibility (e.g. measured as green space acreage) appears adequate (Yasumoto et al. 2021).

This uneven distribution of barriers among groups links to inequalities across the different dimensions of justice (Schlosberg 2009; Loos et al. 2023), although their analysis was not an explicit goal of our study. As in many studies (Guedes et al. 2025), the distributional dimension was most represented, through barriers related to the costs and benefits people derive from nature - such as low availability or quality of providing spaces. The capabilities dimension was also broadly reflected, through barriers such as lack of time, poor health, or financial costs, and often overlapping with distributional inequities. Procedural justice appeared implicitly through participant comments on rights and conflicts over space use - hikers fearing encounters with hunters or guard dogs, hunters asserting their right to hunt, reactions to new access restrictions to formerly open private areas - gesturing at the underlying question of whose use of space is legitimised by governance structures. Fewer barriers were related to the recognitional dimension, specifically whether NCP-providing spaces matched participants’ personal values and cultural identity (“culture”- and “belonging”-related barriers). These were rarely selected, likely reflecting the over-representation of people who are already familiar with these spaces, which per se highlights a limitation in the recognitional dimension of our study. Other groups - such as people from ethnic minorities - might encounter these barriers more frequently but were not explicitly targeted by our sampling design. They may have had limited capacity to learn about and complete the survey, meaning their barriers are likely underestimated.

### Access barriers and nature disconnection impact quality of life

Barriers to access nature can also be understood as forms of nature disconnection (Beery et al., 2023). We mostly identified barriers related to the individual-level dimensions of disconnection, either material (quality, quantity) or cognitive. Conversely, barriers related to sociocultural and institutional disconnection (social insecurity, belonging, cultural barriers) were less prevalent. Such barriers - in particular related to safety and lack of representativeness - disproportionately affect racialised groups (Dennett et al. 2025). However, considering our sampling approach, most respondents were likely to be already interested in, knowledgeable about and/or well-connected to nature.

Despite these limitations, our results showed that the number of barriers reported and nature connectedness, as measured by the Inclusiveness of Nature in Self scale, were driven by similar factors - with a prevalence of the rurality of the respondent’s residence affecting positively nature connectedness (hypothesis H2), and negatively the number of barriers. Both factors (in the case of number of barriers, through satisfaction with NCP) strongly impacted quality of life (supporting H4 and H6). The effect of satisfaction with NCP on quality of life, in particular, was strong - more than that of income, which is generally strongly correlated with quality of life. This might be due to the announced focus of the questionnaire on NCP leading respondents to think of their quality of life through a lens of nature. From our hypothesis H3, we expected a negative effect of the number of barriers on nature connectedness. While the opposite causality (higher nature connectedness lowering barriers, such as fear or lack of knowledge) likely exists too, we did not have the possibility to distinguish between the two pathways and chose the most straightforward direction. Surprisingly, we did not find any correlation between connectedness and the number of barriers reported. While this may reflect the way our survey was framed around NCP rather than time spent in nature, it points to the idea that nature connectedness is more than just a moderator of the relationship between nature contact and quality of life (Martin et al. 2020) - it is also an individual trait linked to greater life satisfaction. While experimental manipulations of these parameters are challenging, they might be needed to untangle their causal relationships.

Nevertheless, our findings suggest that urban residents, women and disadvantaged groups (low-income individuals and people with low physical mobility) tend to experience more barriers, lower connectedness, and lower reported quality of life. These effects likely accumulate alongside other disadvantages, vulnerabilities, and reduced capacity to cope with stressors, ultimately reinforcing inequities. For example, urban residents from the study area - particularly those living in dense, low-income areas - face especially limited access to NCP providing temperature regulation (Neyret et al. 2026). Low income may reduce one’s ability to cope with heatwaves, while poor health and limited mobility may respectively heighten vulnerability to heat and hinder the ability to move to cooler areas (Aznarez et al. 2024). While interactions between our socio-demographic variables were weak (results not presented), these patterns point towards the importance of intersectionality: individuals belonging simultaneously to multiple disadvantaged groups often face especially pronounced barriers to accessing nature and deriving benefits for their well-being (Abdulla et al. 2025; Colley et al. 2022).

### Implications for policy

Our findings show stark disparities in access to nature and their repercussions for quality of life, and highlight the need for environmental and social policies that explicitly recognise and address the unequal distribution of barriers to accessing nature. Addressing these disparities and advancing environmental justice requires integrated policies that acknowledge both the multidimensional nature of barriers to accessing nature and the complex ways these barriers shape human well-being through NCP. For instance, Lejemtel et al. (2025) found that only a small fraction of Parisian policy documents addressed environmental justice in relation to green spaces, and even fewer explicitly targeted vulnerable groups; they also highlighted how sectoral fragmentation hampers the implementation of more integrative strategies.

Current policies pay limited attention to the social, relational, and psychosocial factors that shape who can meaningfully benefit from nature. This limitation is evident in policies that focus on facilitating access to nature or increasing the availability and quality of green spaces. Examples include Sweden’s right of public access (Sandell and Fredman 2010), which grants all individuals the freedom to enter and enjoy natural areas regardless of land ownership. At the European level, the EU Biodiversity Strategy for 2030, the Green Infrastructure Strategy, and the Nature Restoration Law similarly prioritise the expansion and ecological enhancement of green and natural spaces. Urban greening guidance often reinforces this emphasis on quantity and spatial distribution, such as the widely used 3-30-300 rule (Konijnendijk 2023).

There are, nonetheless, emerging signs of policy recognition that access barriers extend beyond spatial provision. Recent initiatives increasingly emphasise the creation of inclusive green spaces tailored to the needs of groups most affected by limited access (PHE 2020), illustrated by efforts such as local initiatives to create therapeutic gardens in Tallinn (Estonia) and Zagreb (Croatia) (EEA 2022) or the development of a network of inclusive parks in Barranquilla (Colombia) (APC Colombia 2025). At larger scales, the SDG 11 (Target 11.7) commits to providing by 2030 universal access to safe, inclusive and accessible green and public spaces, in particular for persons with disabilities (UN 2015). Shifting policy attention from the mere presence of green spaces to the actual ability of diverse populations to engage with and benefit from NCP would allow for more equitable improvements in well-being and support the development of socially just, nature-connected, and climate-resilient cities.

## Conclusion

This study contributes to a growing body of literature on environmental justice and nature access by providing a quantitative comparison of barriers to a wide range of NCP across socio-demographic groups, and by explicitly linking, for the first time, these barriers to nature connectedness and quality of life. Within the validity domain of our study, we identified contrasted responses in terms of the number and type of barriers encountered and their repercussions on quality of life. Disparities might be even more pronounced in the general population. Nature connectedness also contributed to quality of life independently from barriers, highlighting that benefits from nature involve not only physical access but also deeper relational dimensions. The typology of barriers and their relative frequency are likely to vary across socio-cultural contexts; future studies should therefore conduct comparative and intersectional analyses across contexts and social groups, using methodologies that can target underrepresented populations. Finally, establishing the causal direction of the relationships identified here - in particular between barriers, connectedness, and quality of life - would require experimental designs that target specific barriers and observe repercussions on nature connectedness and well-being.

## Acknowledgements

The European Union’s Horizon 2020 research and innovation programme under the Marie Skłodowska-Curie grant agreement No 101104374 provided funding to MN and SL. Project RECONNECT, Biodiversa+ , the European Biodiversity Partnership under the 2021–2022 BiodivProtect joint call for research proposals, provided funding to SL by the ANR Agence Nationale de la Recherche (ANR-22-EBIP-0009-06).

## Data and code availability

The anonymized data to reproduce the analysis has been published Recherche Data Gouv under doi https://doi.org/10.57745/ENKAUD. The script can be found on github: https://github.com/mneyret/Unequal_barriers_AMBIO_2026.

