## Supplementary material for "Unequal barriers to nature’s contributions to people impact quality of life"

### Supplementary results

#### Distribution of respondents across socio-demographic groups

Figures S1 and S2 the distribution of the respondent across socio-demographic factors. As shown in Fig. 1, respondents were relatively well balanced in terms of gender and age, although respondents over 75 years were under represented. Intellectual professions, employees, students and retirees were the most common socio-professional categories. Fig. S2 shows the main axes of variations across the considered factors.

**Figure S1.** Distribution of respondents. Frequency of respondents across age, gender, socio-professional status and childhood environment. NA: missing values (declined to answer).

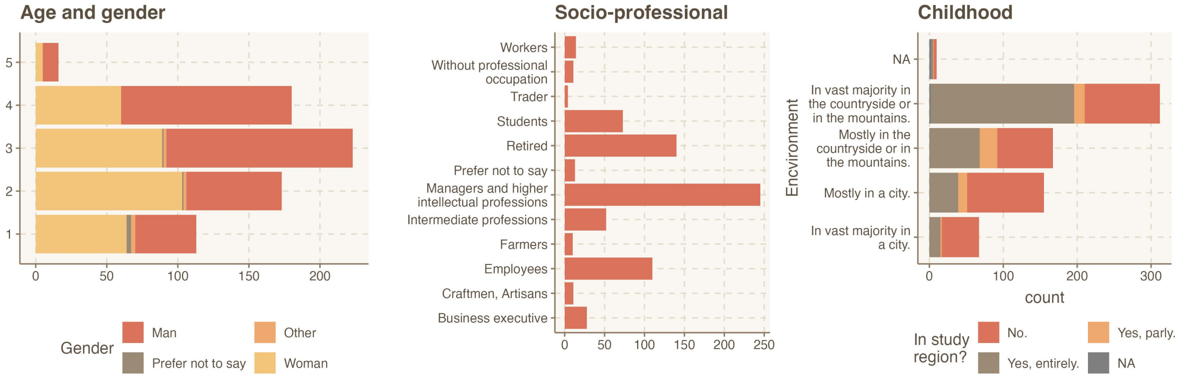

**Figure S2.** PCA of socio-demographic variables. The first axis of variation (Dim 1) is a gradient from urban residents who spent little time in nature and had low nature connectedness, to rural residents with higher nature connectedness, who spent more time in nature, and typically had lived in the region for longer (23.9% variance). The second main axis of variation (Dim 2) is a gradient from low to high income, which was also associated with childhood environment (16.4% ). Axes 3 and 4 are respectively associated with physical mobility (13.8%) and gender (11.1%).
*Diploma: highest diploma; income: annual income, Mobility: self-reported physical mobility, Age: age class, TimeRegion: time spent in the study region (as classes), Rural: degree of rurality of the residence, ChildhoodRegion: to which degree childhood was spent in the region, ChildhoodRural: to which degree childhood was spent in a rural area, Man: gender (1 for men, 0 for women and others). All variables were converted to discrete numerical variables.*

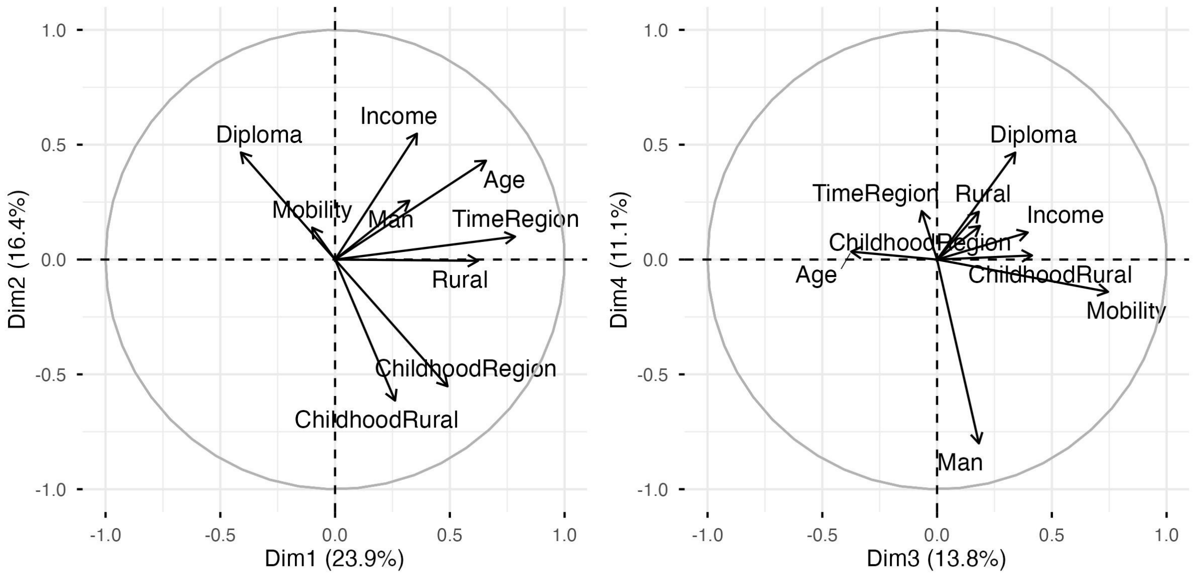

##### Frequency of barriers across NCP and NCP categories

The main barriers differed across NCP categories (Fig. S3). As non-material NCP were the most commonly chosen category (58% of all NCP selected by respondents), the most common barriers for this category were consistent with the overall results, with the two most reported barriers for this category being lack of time (reported for 18% of selected NCP) and crowding (18%). The main barriers reported for regulating NCP (36% of all selected NCP) were low availability (37%) or low ecosystem quality (e.g., degradation or pollution) of green or natural spaces (19%), and lack of knowledge regarding where to find such spaces (18%). The main barriers reported for material NCP (6% of selected NCP) were lack of time (26%), low availability (20%) and high costs (13%). These overall results slightly differed across individual NCP within NCP categories (Figure S4), but were overall consistent. For instance, lack of time and crowding were among the three most common barriers for all non-material NCP, while low quality was a common barrier for aesthetic enjoyment, urban recreation and biodiversity conservation, but not so much for hunting, rural recreation or harvesting.

**Figure S3.** Frequency of barriers reported for each category of NCP. Shaded areas show barriers that were added by the respondents although they were not proposed for this NCP category (i.e., except for temperature regulation in the Regulating NCP panel). Barriers marked with an asterisk (*) were added by the respondents and not present in the initial lists.

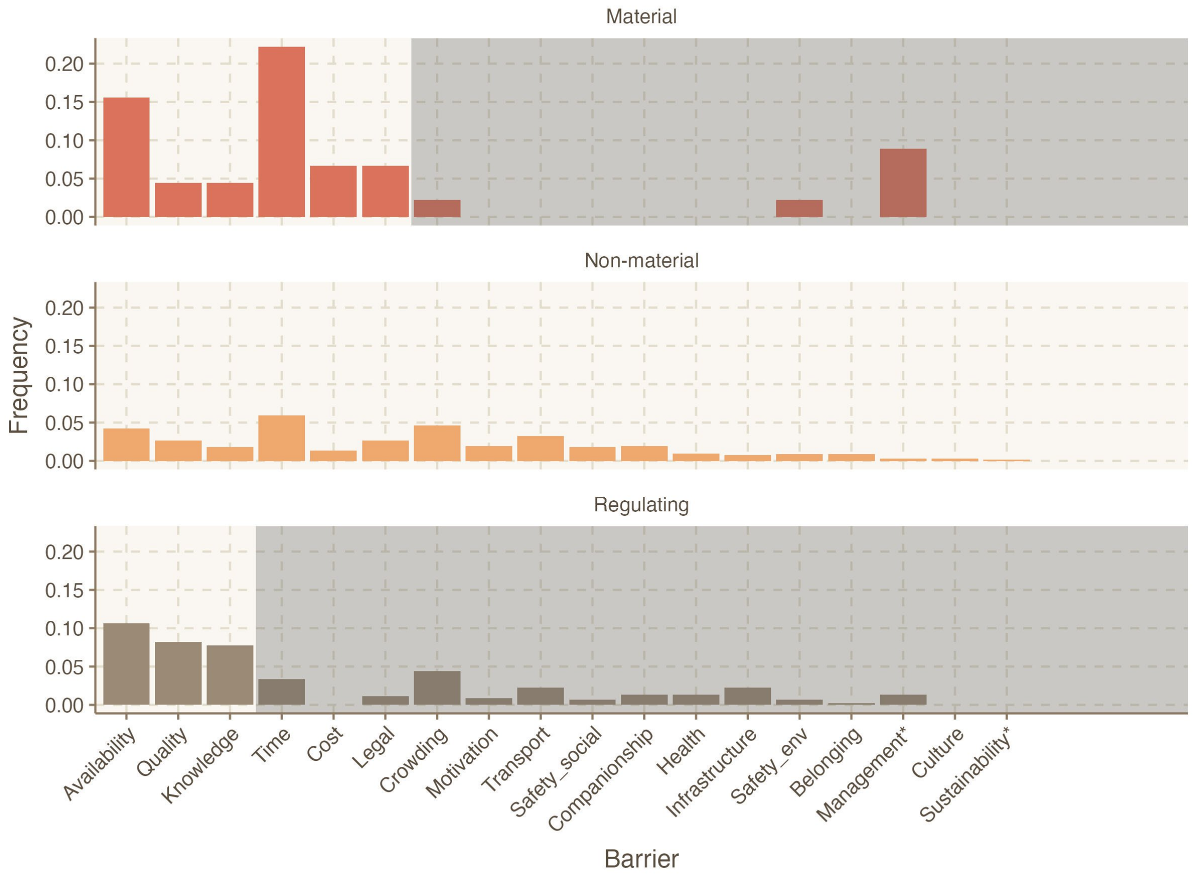

**Figure S4**. Frequency of barriers reported for each NCP. Number of times each barrier was reported, standardised by the number of times the NCP was chosen.

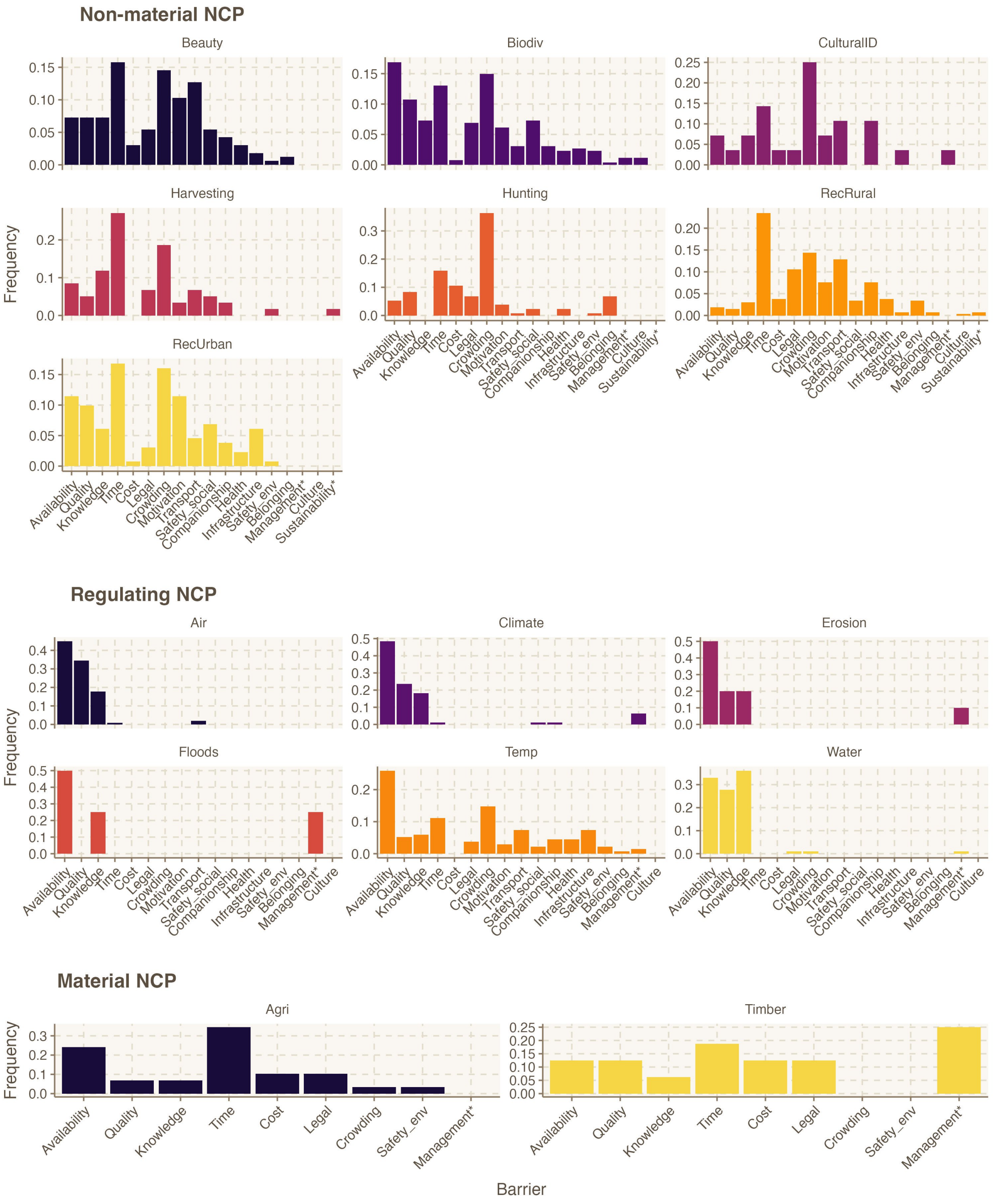

**Table S1.** Model parameters for the impact of socio-demographic variables on the number of barriers reported, by NCP category.

*Income: annual income class; Mobility: self-reported physical mobility; Rural: degree of rurality of the residence; Man: gender (1 for men, 0 for women and others). All variables were converted to discrete numerical variables.*

| NCP category | Explanatory variable | Estimate | Standard error | P-value |
|---|---|---|---|---|
| Material | Income | -0.04 | 0.16 | 0.79 |
| | Man | 0.31 | 0.47 | 0.51 |
| | Mobility | -0.42 | 0.33 | 0.20 |
| | Rural | -0.06 | 0.08 | 0.48 |
| Non-material | Income | **-0.12** | **0.04** | **0.002**** |
| | Man | -0.13 | 0.12 | 0.25 |
| | Mobility | **-0.22** | **0.08** | **0.009**** |
| | Rural | **-0.09** | **0.02** | **<.001***** |
| Regulating | Income | -0.03 | 0.04 | 0.41 |
| | Man | **-0.28** | **0.13** | **0.029*** |
| | Mobility | -0.05 | 0.09 | 0.55 |
| | Rural | **-0.09** | **0.03** | **<.001***** |

**Table S2.** Model parameters for the impact of socio-demographic variables on the likelihood of reporting each individual barrier. Models for each barrier were run independently.
*Income: annual income class; Mobility: self-reported physical mobility; Rural: degree of rurality of the residence; Man: gender (1 for men, 0 for women and others). All variables were converted to discrete numerical variables.*

| Barrier | Explanatory variable | Estimate | Standardised error | P-value |
|---|---|---|---|---|
| Availability | Income | -0.02 | 0.05 | 0.70 |
| | Man | 0.08 | 0.16 | 0.62 |
| | Mobility | -0.20 | 0.11 | 0.076 |
| | **Rural** | **-0.14** | **0.04** | **<.001*****|
| Companionship | Income | -0.26 | 0.18 | 0.15 |
| | Man | -0.57 | 0.56 | 0.30 |
| | Mobility | -0.41 | 0.35 | 0.24 |
| | Rural | -0.17 | 0.12 | 0.15 |
| Cost | **Income** | **-0.71** | **0.20** | **<.001*****|
| | Man | -0.10 | 0.51 | 0.85 |
| | Mobility | 0.47 | 0.42 | 0.26 |
| | Rural | 0.02 | 0.10 | 0.81 |
| Crowding | Income | -0.09 | 0.08 | 0.27 |
| | Man | 0.24 | 0.23 | 0.29 |
| | Mobility | -0.22 | 0.16 | 0.18 |
| | Rural | -0.05 | 0.05 | 0.27 |
| Health | Income | 0.01 | 0.13 | 0.92 |
| | Man | -2.10 | 0.87 | 0.16 |
| | **Mobility** | **-1.52** | **0.20** | **<.001*****|
| | Rural | 0.06 | 0.08 | 0.40 |
| Infrastructure | Income | -0.01 | 0.30 | 0.98 |
| | Man | -0.74 | 1.01 | 0.46 |
| | Mobility | -0.12 | 0.67 | 0.86 |

| | | | | |
|---|---|---|---|---|
| | Rural | -0.23 | 0.22 | 0.30 |
| Knowledge | Income | -0.04 | 0.08 | 0.61 |
| | **Man** | **-0.77** | **0.26** | **0.003**** |
| | Mobility | -0.11 | 0.17 | 0.52 |
| | **Rural** | **-0.15** | **0.05** | **0.004**** |
| Legal | Income | -0.20 | 0.16 | 0.23 |
| | Man | 0.62 | 0.45 | 0.17 |
| | Mobility | 0.20 | 0.36 | 0.58 |
| | Rural | 0.06 | 0.10 | 0.55 |
| Management | Income | 0.00 | 0.37 | 1 |
| | Man | 0.00 | 0.70 | 1 |
| | Mobility | 0.00 | 0.56 | 1 |
| | Rural | 0.00 | 0.13 | 1 |
| Motivation | Income | -0.15 | 0.13 | 0.24 |
| | Man | -0.26 | 0.40 | 0.51 |
| | **Mobility** | **-0.59** | **0.27** | **0.030*** |
| | **Rural** | **-0.20** | **0.09** | **0.026*** |
| Quality | **Income** | **-0.21** | **0.08** | **0.005**** |
| | Man | 0.09 | 0.22 | 0.67 |
| | Mobility | 0.10 | 0.16 | 0.53 |
| | Rural | -0.05 | 0.05 | 0.29 |
| Safety (environment) | Income | -0.17 | 0.37 | 0.65 |
| | Man | -1.24 | 1.39 | 0.37 |
| | Mobility | 0.35 | 0.88 | 0.69 |
| | Rural | 0.07 | 0.22 | 0.75 |
| Safety (social) | Income | -0.02 | 0.24 | 0.94 |
| | Man | -0.83 | 0.81 | 0.30 |
| | Mobility | -0.19 | 0.52 | 0.71 |
| | Rural | 0.12 | 0.15 | 0.43 |

| | | | | |
|---|---|---|---|---|
| Time | Income | 0.03 | 0.07 | 0.70 |
| | Man | -0.15 | 0.21 | 0.46 |
| | Mobility | -0.05 | 0.16 | 0.77 |
| | **Rural** | **-0.15** | **0.05** | **0.001*****|
| Transport | Income | -0.13 | 0.11 | 0.25 |
| | **Man** | **-1.42** | **0.42** | **<.001*****|
| | Mobility | -0.11 | 0.24 | 0.66 |
| | **Rural** | **-0.47** | **0.11** | **<.001*****|

**Table S3.** Impact of socio-demographic variables on satisfaction with the NCP provided, nature connectedness, and quality of life. Results from Cumulative Link Mixed Models. *To run the model for quality of life, it was necessary to merge the two lowest categories into one factor due to the very low number of responses for the lowest category.
*Income: annual income class; Mobility: self-reported physical mobility; Rural: degree of rurality of the residence; Man: gender (1 for men, 0 for women and others). All variables were converted to discrete numerical variables.*

| Response variable | Explanatory variable | Estimate | Std. Error | z value | Pr(>\|z\|) |
|---|---|---|---|---|---|
| Satisfaction | **Gender (man)** | **0.06** | **0.03** | **2.45** | **0.01 *** |
| | **Mobility** | **0.37** | **0.09** | **4.12** | **<.001 ***** |
| | Income | 0.08 | 0.12 | 0.65 | 0.51 |
| | Rural | 0.08 | 0.04 | 1.90 | 0.06 |
| Quality of life* | Gender (man) | 0.02 | 0.12 | 0.14 | 0.89 |
| | **Mobility** | **0.54** | **0.09** | **5.94** | **<.001 ***** |
| | **Income** | **0.23** | **0.04** | **5.55** | **<.001 ***** |
| | **Rural** | **0.17** | **0.03** | **7.00** | **<.001 ***** |
| Nature connectedness | **Gender (man)** | **0.25** | **0.11** | **.0.11** | **0.004 **** |
| | **Mobility** | **0.46** | **0.09** | **5.34** | **<.001 ***** |
| | Income | 0.04 | 0.04 | 1.02 | 0.31 |
| | **Rural** | **0.25** | **0.03** | **10.07** | **<.001 ***** |

**Table S4.** SEM results. Model fit indices: pvalue = 0.230, cfi = 0.999, rmsea = 0.018.
*Income: annual income class; Mobility: self-reported physical mobility; Rural: degree of rurality of the residence; Man: gender (1 for men, 0 for women and others); QoL: quality of life; Satisfaction: satisfaction with the NCP provided; Connectedness: nature connectedness. All variables were converted to discrete numerical variables.*

| Path | Estimate | Standard error | P-value | Confidence interval |
|---|---|---|---|---|
| **Satisfaction ~ number of barriers** | **-0.17***** | **0.02** | **0.00** | **[-0.22, -0.13]** |

| | | | | |
|---|---|---|---|---|
| **Satisfaction ~ Mobility** | **0.16*** ** | **0.04** | **0.00** | **[0.09, 0.23]** |
| Satisfaction ~ Man | -0.04 | 0.05 | 0.37 | [-0.13, 0.05] |
| Satisfaction ~ Income | 0.02 | 0.02 | 0.20 | [-0.01, 0.05] |
| Satisfaction ~ Rural | 0.00 | 0.01 | 0.95 | [-0.02, 0.02] |
| **QoL ~ Satisfaction** | **0.60*** ** | **0.12** | **0.00** | **[0.36, 0.83]** |
| QoL ~ Mobility | 0.03 | 0.04 | 0.42 | [-0.05, 0.12] |
| QoL ~ Man | -0.00 | 0.04 | 0.98 | [-0.08, 0.08] |
| **QoL ~ Income** | **0.05** ** | **0.02** | **0.00** | **[0.02, 0.08]** |
| **QoL ~ Rural** | **0.03** ** | **0.01** | **0.00** | **[0.01, 0.05]** |
| **QoL ~ Connectedness** | **0.09*** ** | **0.02** | **0.00** | **[0.05, 0.13]** |
| **number of barriers ~ Mobility** | **-0.14** ** | **0.05** | **0.01** | **[-0.24, -0.04]** |
| **number of barriers ~ Man** | **-0.18** ** | **0.06** | **0.00** | **[-0.30, -0.07]** |
| **number of barriers ~ Income** | **-0.09*** ** | **0.02** | **0.00** | **[-0.13, -0.04]** |
| **number of barriers ~ Rural** | **-0.08*** ** | **0.01** | **0.00** | **[-0.11, -0.06]** |
| **Connectedness ~ Man** | **0.15*** ** | **0.04** | **0.00** | **[0.07, 0.23]** |
| Connectedness ~ Income | 0.01 | 0.01 | 0.71 | [-0.02, 0.03] |
| **Connectedness ~ Rural** | **0.10*** ** | **0.01** | **0.00** | **[0.08, 0.12]** |
| **Connectedness ~ Mobility** | **0.14*** ** | **0.03** | **0.00** | **[0.08, 0.21]** |
| Connectedness ~ number of barriers | -0.02 | 0.02 | 0.34 | [-0.06, 0.02] |
| **Satisfaction ~~ QoL** | **-0.29*** ** | **0.08** | **0.00** | **[-0.44, -0.14]** |
| **Mobility ~~ Man** | **0.03*** ** | **0.01** | **0.00** | **[0.02, 0.05]** |

| | | | | |
|---|---|---|---|---|
| **Mobility ~~ Income** | **0.09*** ** | **0.03** | **0.00** | **[0.04, 0.15]** |
| **Man ~~ Income** | **0.12*** ** | **0.02** | **0.00** | **[0.09, 0.16]** |
| Man ~~ Rural | 0.06 | 0.03 | 0.06 | [-0.00, 0.12] |
| **Income ~~ Rural** | **0.72*** ** | **0.09** | **0.00** | **[0.54, 0.90]** |
| **Variance:Satisfaction** | **0.61*** ** | **0.02** | **0.00** | **[0.56, 0.65]** |

**Table S5.** Results from alternative SEM modelling using piecewise SEM. Model fit was poor: Chi-Squared = 179.015 (P-value = 0); Fisher's C = 4.811 (P-value = 0.09).
*Income: annual income class; Mobility: self-reported physical mobility; Rural: degree of rurality of the residence; Man: gender (1 for men, 0 for women and others); QoL: quality of life; Satisfaction: satisfaction with the NCP provided; Connectedness: nature connectedness. All variables were converted to discrete numerical variables.*

| Response | Predictor | Standard Estimate | Std.Error | DF | Crit.Value | P.Value |
|---|---|---|---|---|---|---|
| **Satisfaction** | **nbarriers** | **-0.29** | **0.02** | **1073** | **144.82** | **<.001 *** ** |
| **Satisfaction** | **Mobility** | **0.09** | **0.03** | **1050** | **14.92** | **<.001 *** ** |
| Satisfaction | Man | -0.02 | 0.04 | 1043 | 0.76 | 0.3839 |
| Satisfaction | Income | 0.0232 | 0.01 | 979 | 1.01 | 0.3155 |
| Satisfaction | Rural | 0.02 | 0.01 | 1001 | 0.76 | 0.3851 |
| **QV** | **nbarriers** | **-0.13** | **0.02** | **599** | **27.34** | **<.001 *** ** |
| **QV** | **Satisfaction** | **0.15** | **0.02** | **91** | **32.87** | **<.001 *** ** |
| **QV** | **Mobility** | **0.10** | **0.03** | **1038** | **18.07** | **<.001 *** ** |
| QV | Man | -0.01 | 0.03 | 231 | 0.38 | 0.5398 |
| **QV** | **Income** | **0.13** | **0.01** | **974** | **27.81** | **<.001 *** ** |

| | | | | | | |
|---|---|---|---|---|---|---|
| **QV** | **Rural** | **0.11** | **0.01** | **694** | **18.06** | **<.001 *** ** |
| **QV** | **Connectedness** | **0.11** | **0.02** | **818** | **16.36** | **<.001 *** ** |
| **nbarriers** | **Mobility** | **-0.08** | **0.05** | **1552** | **-3.49** | **<.001 *** ** |
| **nbarriers** | **Man** | **-0.08** | **0.07** | **1552** | **-3.07** | **0.002 ** ** |
| **nbarriers** | **Income** | **-0.09** | **0.02** | **1552** | **-3.67** | **<.001 *** ** |
| **nbarriers** | **Rural** | **-0.18** | **0.01** | **1552** | **-6.72** | **<.001 *** ** |
| **Connectedness** | **Man** | **0.07** | **0.04** | **1049** | **8.99** | **0.003 ** ** |
| Connectedness | Income | 0.01 | 0.01 | 1052 | 0.34 | 0.56 |
| **Connectedness** | **Rural** | **0.09** | **0.01** | **1056** | **123.79** | **<.001 *** ** |
| **Connectedness** | **Mobility** | **0.16** | **0.03** | **1118** | **31.84** | **<.001 *** ** |
| Connectedness | nbarriers | -0.02 | 0.02 | 1120 | 1.51 | 0.22 |

**Table S6.** Results from alternative SEM model (as shown in Table S4) but without using different weights for hunters. Model fit was poor: Chi-Squared = 179.015 (P-value = 0); Fisher's C = 4.811 (P-value = 0.09).

*Income: annual income class; Mobility: self-reported physical mobility; Rural: degree of rurality of the residence; Man: gender (1 for men, 0 for women and others); QoL: quality of life; Satisfaction: satisfaction with the NCP provided; Connectedness: nature connectedness. All variables were converted to discrete numerical variables.*

| Path | Estimate | Standard error | P-value | Confidence interval |
|---|---|---|---|---|
| **Satisfaction ~ nbarriers** | **-0.20*** ** | **0.02** | **0.00** | **[-0.24, -0.17]** |
| **Satisfaction ~ Mobility** | **0.14*** ** | **0.03** | **0.00** | **[0.07, 0.20]** |
| Satisfaction ~ Man | -0.02 | 0.04 | 0.67 | [-0.10, 0.06] |
| Satisfaction ~ Income | 0.01 | 0.01 | 0.71 | [-0.02, 0.03] |

| | | | | |
|---|---|---|---|---|
| Satisfaction ~ Rural | 0.01 | 0.01 | 0.53 | [-0.01, 0.02] |
| **QoL ~ satisfaction** | **0.48*** ** | **0.08** | **0.00** | **[0.32, 0.64]** |
| QoL ~ Mobility | 0.06 | 0.03 | 0.07 | [-0.00, 0.12] |
| QoL ~ Man | -0.02 | 0.04 | 0.66 | [-0.09, 0.05] |
| **QoL ~ Income** | **0.05*** ** | **0.01** | **0.00** | **[0.03, 0.08]** |
| **QoL ~ Rural** | **0.03*** ** | **0.01** | **0.00** | **[0.02, 0.05]** |
| **QoL ~ Connectedness** | **0.09*** ** | **0.02** | **0.00** | **[0.05, 0.13]** |
| **nbarriers ~ Mobility** | **-0.15*** ** | **0.04** | **0.00** | **[-0.23, -0.06]** |
| **nbarriers ~ Man** | **-0.14* ** | **0.05** | **0.01** | **[-0.25, -0.03]** |
| **nbarriers ~ Income** | **-0.07*** ** | **0.02** | **0.00** | **[-0.11, -0.03]** |
| **nbarriers ~ Rural** | **-0.08*** ** | **0.01** | **0.00** | **[-0.10, -0.05]** |
| **Connectedness ~ Man** | **0.20*** ** | **0.04** | **0.00** | **[0.12, 0.27]** |
| Connectedness ~ Income | -0.02 | 0.01 | 0.16 | [-0.05, 0.01] |
| **Connectedness ~ Rural** | **0.10*** ** | **0.01** | **0.00** | **[0.09, 0.12]** |
| **Connectedness ~ Mobility** | **0.14*** ** | **0.03** | **0.00** | **[0.08, 0.20]** |
| Connectedness ~ nbarriers | -0.03 | 0.02 | 0.06 | [-0.07, 0.00] |
| **Satisfaction ~~ QoL** | **-0.21*** ** | **0.05** | **0.00** | **[-0.31, -0.11]** |
| **Mobility ~~ Man** | **0.02** ** | **0.01** | **0.00** | **[0.01, 0.04]** |
| **Mobility ~~ Income** | **0.11*** ** | **0.02** | **0.00** | **[0.06, 0.15]** |
| **Man ~~ Income** | **0.14*** ** | **0.02** | **0.00** | **[0.10, 0.18]** |
| **Man ~~ Rural** | **0.22*** ** | **0.03** | **0.00** | **[0.16, 0.28]** |

| | | | | |
|---|---|---|---|---|
| **Income ~~ Rural** | **0.69*** ** | **0.09** | **0.00** | **[0.52, 0.87]** |
| **Variance:satisfaction** | **0.60*** ** | **0.02** | **0.00** | **[0.56, 0.64]** |

#### Supplementary methods

##### SEM models

We fitted a Structural Equation Model (SEM) linking socio-demographic variables and quality of life, with the number of barriers reported, satisfaction with NCP, and nature connectedness as mediating factors. We fitted the SEM using two approaches.

- First using a traditional SEM (package lavaan, R). Due to model limitations, each path was considered linear and no random effect was included. However, residual covariations between exogenous variables (gender, mobility, income, rurality) could be included. The resulting model fitted correctly the data (pvalue = 0.230, cfi =0.99, rmsea =0.018).
- We also fitted the same model using a piecewise approach (piecewiseSEM package). Models with ordinal variables as response variables (satisfaction, quality of life, nature connectedness) were modeled as linear mixed effects with NCP as a random factor because it was not possible to use cumulative link models in piecewiseSEM to account for the ordinal characteristics of the variable. The number of barriers was modelled using generalized mixed models with a poisson family. Because residual covariations between exogenous variables could not be included in piecewiseSEM, the model fit was poor (Chi-Squared = 200.6 with P-value = 0; Fisher's C = 35.4 with P-value = 0.09).

Despite their limitations, both modelling options yielded similar results. We present the results of the first approach in the main text (model estimates detailed in Table S4), and also show the results of the piecewise SEM in Table S5.

##### Survey

We reproduce here the parts of the online survey. Sections or questions irrelevant to the present study are omitted.

***Nature, Green Spaces, and Quality of Life in the Greater Grenoble Region***

[This study] aims to analyze the relationships between people and nature, and to examine disparities in access to natural areas and green spaces in the greater Grenoble region (green zone on the map opposite, extending roughly from Chambéry to Vienne, down to the https://doi.org/10.1007/s13280-026-02482-z

south of Trièves). Here, “nature” refers to living beings (animals, plants), ecosystems, or landscapes that make up our region. “Green spaces” include all areas with grass, bushes, and/or trees, such as parks, private gardens, or community gardens. The survey focuses on people’s relationships with nature throughout the year, across all seasons. [...]

- First, please indicate where you live. [Map question]

- We will now ask you a few questions related to your quality of life and well-being. Please indicate whether you agree with the following statements:

  I am satisfied with my quality of life and well-being.

  - Strongly disagree
  - Somewhat disagree
  - Somewhat agree
  - Strongly agree

Comments (optional):

**Priorities, satisfaction and barriers related to NCP**

- We are interested in the different benefits provided by nature in the study area and how important they are to you. For each of the following benefits, please indicate its importance for your quality of life and well-being, across all seasons (summer, winter, etc.). To do this, you may distribute up to **30 points in total** among the different benefits.
  - **Beauty of nature and landscapes**: flowers (wild or planted), trees, rivers, mountains…
  - **Habitats for biodiversity**: creation and maintenance of high-quality habitats for biodiversity (plants, animals…).
  - **Cooling of temperature**: shade and freshness provided by vegetation (e.g., it feels cooler near trees or grass compared to concrete).
  - **Flood protection**: vegetation protecting against floods, for example by improving water infiltration into soil.
  - **Outdoor leisure in town or nearby**: walking, sports, relaxing in parks, private or public gardens, green spaces.
  - **Cultural identity**: species and landscapes that are part of the cultural heritage or identity of the region (e.g., edelweiss is often seen as emblematic of alpine areas).
  - **Water quality**: maintaining water quality in rivers, lakes, and groundwater (e.g., riparian vegetation reducing agricultural pollution).
  - **Global climate regulation**: limiting greenhouse gases in the atmosphere to slow climate change (e.g., forests storing carbon).
  - **Wood production**: local production of wood for construction or heating.
  - **Air quality**: clean, oxygenated, depolluted air thanks to vegetation.

- **Crops and livestock**: local agricultural production of cereals, fruits, vegetables; livestock for milk and meat (cows, sheep…).
- **Erosion and landslide protection**: preventing soil erosion, rockfalls, or landslides (e.g., protective forests reducing risks near roads or housing).
- **Outdoor leisure in the countryside or mountains**: camping, swimming, hiking, walking, relaxing, observing plants and animals.
- **Foraging**: collecting mushrooms, berries, wild plants.
- **Hunting and fishing**: fishing in rivers and lakes, hunting game.

*[For each up to three NCP they gave the most points to, respondents were then requested to answer the following questions. An example is given for landscape aesthetic value; listed barriers differed across NCP as shown in Table 1]*

- I am satisfied with the beauty of the green or natural spaces around me.
    - Strongly disagree
    - Somewhat disagree
    - Somewhat agree
    - Strongly agree
    - I don't know

- If one or more of the following factors prevents you (totally or partially) from accessing and enjoying this benefit, please check the corresponding box. Otherwise, you may skip to the next question.
    - There are no (or not enough) nearby green or natural spaces providing this benefit.
    - The spaces providing this benefit are of poor quality or in bad condition (dirty, degraded…).
    - It's too expensive.
    - I don't know where to go, or which spaces provide this benefit.
    - I am legally not allowed to enjoy this benefit where I would like to (private land, restricted/protected areas).
    - I don't have the time, I'm too busy.
    - There are too many other people, or other people disturb me (noise, crowds…).
    - I cannot easily access the spaces providing this benefit (lack of roads, public transport).
    - I have no one to go with.
    - I don't feel comfortable because I feel isolated or marginalized in these spaces.
    - I don't feel comfortable because the spaces do not fit with my beliefs or culture.
    - There is insufficient infrastructure (benches, toilets, signs…) in these spaces.
    - I lack motivation.
    - I don't feel safe in these spaces because of other people.

- I don't feel safe in these spaces because of the environment (fear of getting lost, falling, tick or insect bites…).
- My health prevents me from accessing or enjoying these spaces (physical condition, pain, allergies, asthma…).
- Other (please specify):

**Personal Background**

Thank you for answering up to this point! We will now ask you a few questions about your personal context.

- **What gender do you identify with?**
  - Woman
  - Man
  - Other
  - Prefer not to say

- **What is your age?**
  - 18–29 years
  - 30–44 years
  - 45–59 years
  - 60–74 years
  - 75 years and over
  - Prefer not to say

- **What is your highest level of education?**
  - No diploma
  - Middle school certificate (Brevet)
  - Vocational qualification (CAP-BEP)
  - High school diploma (Baccalauréat)
  - 2-year postsecondary diploma (Bac +2)
  - 3–5 year university degree or higher (Bac +3, Bac +5 or more)
  - Prefer not to say

- **Town**

- **Postal code**

- **What is your nationality?** [Drop-down menu]

- **How would you describe your physical mobility (ease of walking, moving, exercising…)?**

- Very good
- Good
- Limited
- Very limited
- Prefer not to say

- **Where did you spend your childhood?**
  - Mostly in the countryside or mountains
  - Rather in the countryside or mountains
  - Rather in the city
  - Mostly in the city

- **How long have you lived in the study region?**
  - I don't currently live in the region
  - 5 years or less
  - 5–10 years
  - 10–20 years
  - 20 years or more

- **Did you spend your childhood in the study region?**
  - Yes, entirely
  - Yes, partly
  - No

- **How much time per week (including weekends) do you usually spend in nature or green spaces?**
  - Less than 1 hour
  - More than 1 hour but less than 3 hours
  - More than 3 hours but less than 5 hours
  - More than 5 hours but less than 7 hours
  - More than 7 hours but less than 9 hours
  - More than 9 hours but less than 11 hours
  - 11 hours or more

- **Choose the image that best describes your relationship with nature in the study region.**

- **How many people live in your household (including yourself)?**

- **What is your main socio-professional category?**
  - Farmers
  - Craftspeople
  - Shopkeepers
  - Business owners

    - Executives and higher intellectual professions
    - Intermediate professions
    - Employees
    - Workers
    - Retirees
    - Students
    - Unemployed
    - Prefer not to say

- **In your profession, are you involved in the management of nature, land, green spaces, or the environment?**
    - No
    - Agriculture
    - Livestock
    - Forestry
    - Biodiversity conservation
    - Land-use planning
    - Water management
    - Ecotourism
    - Outdoor sports activities
    - Research
    - Other (please specify):

- **Outside your profession (volunteering, associations…), are you involved in the management of nature, land, green spaces, or the environment?**
    - No
    - Agriculture
    - Livestock
    - Forestry
    - Biodiversity conservation
    - Land-use planning
    - Water management
    - Ecotourism
    - Outdoor sports activities
    - Research
    - Other (please specify):

- **What is your household's gross annual income (salary, rental income, social benefits…)?**
    - €11,000 or less per year
    - €11,001 – €26,000
    - €26,001 – €41,000
    - €41,001 – €56,000
    - €56,001 – €71,000
    - €71,001 or more
    - Prefer not to say
